# Optimizing optogenetic stimulation protocols in auditory corticofugal neurons based on closed-loop spike feedback

**DOI:** 10.1101/505214

**Authors:** Charles-Henri Vila, Ross S Williamson, Kenneth E Hancock, Daniel B Polley

## Abstract

Optogenetics provides a means to probe functional connections between brain areas. By activating a set of presynaptic neurons and recording the activity from a downstream brain area, one can establish the sign and strength of a feedforward connection. One challenge is that there are virtually limitless patterns that can be used to stimulate a presynaptic brain area. Functional influences on downstream brain areas can depend not just on *whether* presynaptic neurons were activated, but *how* they were activated. Corticofugal axons from the auditory cortex (ACtx) heavily innervate the auditory tectum, the inferior colliculus (IC). Despite the anatomical weight of this connection, optogenetic activation of ACtx neurons produced only modest changes in the IC neuron firing rates. To determine whether different modes of cortical activation could more faithfully reveal the strength of feedforward connectivity, we employed a closed-loop evolutionary optimization procedure that tailored voltage command signals to the laser based on firing rate variations recorded from single units in the IC of awake male and female mice. Within minutes, the evolutionary search procedure converged on ACtx stimulation configurations that produced more effective and widespread enhancement of IC unit activity than generic activation parameters. Cortical modulation of midbrain spiking was bi-directional, as the evolutionary search procedure could be programmed to converge on activation patterns that suppressed or enhanced sound-evoked IC firing rate. These findings demonstrate that the feedforward influence between brain areas can vary both in sign and degree depending on how presynaptic neurons are activated in time.

**Significance Statement:** Neurons in deep layers of the auditory cortex (ACtx) make extensive projections to subcortical auditory areas, yet little is known about how these descending projections modulate subcortical sound processing in real time. Here, we leveraged recent advances in multi-channel electrophysiology and optogenetics to record from multiple regions of the inferior colliculus (IC) while optogenetically stimulating cortical neurons expressing Chronos, an ultra-fast channelrhodopsin. To identify ACtx activation patterns associated with the strongest effects on IC firing rates, we applied a machine learning algorithm that utilized the firing rate of single IC neurons to iteratively tailor the voltage command signal sent to the laser. We show that the temporal patterning of ACtx spiking strongly impacts the cortical influence on midbrain sound processing.

## Introduction

Descending projections directly connect the auditory cortex (ACtx) to downstream neurons in the basal ganglia, amygdala and nearly all levels of subcortical auditory processing (Winer, 2006). One major descending projection originates in layers 5 of the ACtx and projects to the inferior colliculus (IC) in the midbrain (Diamond et al., 1969; Beyerl, 1978). These corticocollicular (CCol) projections are glutamatergic, largely ipsilateral, and arise from all areas of the ACtx (Kaneko et al., 1987; Feliciano and Potashner, 1995; Winer et al., 1998; Coomes et al., 2005). Although CCol projections primarily target the external and dorsal cortex of the IC, sparse CCol axon collaterals are also found in the central nucleus, which, when combined with dense intracollicular connections suggest that the descending CCol projections could potentially modulate neurons in all regions of the IC, either through direct projections or local polysynaptic connections (Ito et al., 2016).

Numerous studies have investigated the role of corticocollicular projections in auditory processing by focally stimulating or reversibly cooling the ACtx and characterizing the effect on downstream IC responses. Short-term activation or inactivation of ACtx can modify auditory tuning to various dimensions of sound frequency and time (Ma and Suga, 2001a, 2001b; Yan and Ehret, 2002; Yan and Zhang, 2005; Zhou and Jen, 2005; Nakamoto et al., 2008; Robinson et al., 2016), where the direction of change depends upon the topographic alignment of ACtx and IC neurons (Ma and Suga, 2001b; Yan and Ehret, 2002). Although these results suggest that descending feedback plays a substantial role in auditory processing, interpretation is challenging for multiple reasons. Focal microstimulation or cooling indiscriminately manipulates many classes of ACtx neurons, including subcerebral and intratelencephalic projection neurons, interneurons, thalamic axon terminals, and even axons of passage. This limitation is further compounded by the use of anesthetized preparations, which disproportionately affect efferent projections systems (Chambers et al., 2012). These conventional approaches have been fruitful for studies of short- and long-term plasticity processes that unfold in the IC when the ACtx is continuously microstimulated, cooled, or inhibited for periods lasting several minutes to many days. Less is known however, about the contribution of the ACtx to real time modulation of sound processing in downstream target neurons.

Here, we leverage advances in multi-channel electrophysiology and the development of ultra-sensitive opsins to record from multiple areas of the IC while stimulating the ACtx with different temporal patterns of light. We show that using a closed-loop adaptive algorithm to optimize patterns of ACtx activation can lead to either enhancement or suppression of downstream neurons in the IC, demonstrating that the effect of corticofugal feedback can differ dependent on how presynaptic neurons are activated in time.

## Materials & Methods

### Mice

All procedures were approved by the Massachusetts Eye and Ear Infirmary Animal Care and Use Committee and followed the guidelines established by the National Institute of Health for the care and use of laboratory animals. All procedures were performed on 10 CBA/CaJ mice of either sex. Mice were maintained under a regular light cycle (light: 7am - 7pm, dark: 7pm - 7am) with ad libitum access to food and water.

### Surgical Procedures

#### Virus-mediated gene delivery

Mice aged 6-8 weeks were anesthetized using 1-2% isoflurane in oxygen. A homoeothermic blanket system was used to maintain core body temperature at approximately 36.5°C (Fine Science Tools). The surgical area was first shaved and prepared with iodine and ethanol before being numbed with a subcutaneous injection of lidocaine (5 mg/ml). An incision was made to the right side of the scalp to expose the skull overlying the ACtx. The temporalis muscle was then retracted and 2 burr holes were made along the suture line where the temporalis muscle attaches to the skull, approximately 1.5 - 2.5 mm rostral to the lambdoid suture. A motorized stereotaxic injector (Stoelting Co.) was used to inject 0.5 µl of AAV2/8-Synapsin-Chronos-GFP into each burr hole approximately 500 µm below the pial surface with an injection rate of 0.05-0.1 µl/min. Following the injection, the surgical area was sutured, antibiotic ointment was applied to the wound margin, and an analgesic was administered (Buprenex, 0.05 mg/ kg). Neurophysiology experiments began 3-4 weeks following virus injection.

#### Preparation for awake, head-fixed recordings

Mice were brought to a surgical plane of anesthesia, as described above. The dorsal surface of the skull was exposed, and the periosteum was thoroughly removed. The skull was then prepared with 70% ethanol and etchant (C & B Metabond) to ensure an adequate surface for cement application. A custom titanium head plate (eMachineShop) was then cemented to the skull, centered on bregma. For optogenetic stimulation, a multimode optic fiber (0.2 mm core fiber diameter) was either implanted atop the surface of the auditory cortex at the virus injection site (N=5 mice) or 3.5-4 mm below the brain surface to target the fasciculated corticocollicular axon bundle where it first enters the midbrain (N=5 mice). After recovery, all mice were housed individually.

Before the first recording session, mice were briefly anesthetized with isoflurane (1%) while a craniotomy was made atop the IC (1×1 mm centered 0.25 mm caudal to the lambdoid suture, 1 mm lateral to midline). A small chamber was built around the craniotomy with UV-cured cement and filled with lubricating ointment (Bacitracin). At the end of each recording session, the chamber was flushed, filled with fresh ointment, and capped with UV-cured cement (Flow-It ALC). The chamber was removed and rebuilt under isoflurane anesthesia before each subsequent recording session. Typically, 3-5 recording sessions were performed on each animal over the course of one week.

### Neurophysiology

#### Awake, head-fixed preparation

On the day of recording, the head was immobilized by attaching the head plate to a rigid clamp (Altechna). Mice could walk freely on a disk that was mounted atop a low-friction silent rotor and a high-sensitivity optical rotary encoder. Continuous monitoring of the eye and disk rotation confirmed that all recordings were made in the awake condition. Recordings were performed inside a dimly lighted single-wall sound attenuating chamber (Acoustic Systems). Mice were first habituated to head restraint during two 1-hour sessions prior to the first day of recording. Acoustic stimuli were presented via a freefield electrostatic speaker positioned 10 cm from the left ear canal (Tucker-Davis Technologies). Stimuli were calibrated before recording using a wide-band ultrasonic acoustic sensor (Knowles Acoustics, model SPM0204UD5).

#### Data acquisition

At the beginning of each session, a 32-channel, 4-shank, silicon probe (NeuroNexus A4×8-5mm-100-200-177-Z32) was inserted into the IC craniotomy perpendicular to the brain surface using a micromanipulator (Narishige) and a hydraulic microdrive (FHC). Once inserted, the brain was allowed 10-20 minutes to settle before recording began. Raw signals were digitized at 32-bit, 24.4 kHz and stored in binary format (PZS Neurodigitizer and RZS BioAmp Processor; Tucker-Davis Technologies). To minimize artifacts, the common mode signal (channel-averaged neural traces) was subtracted from all channels (Ludwig et al., 2009). Electrical signals were notch filtered at 60 Hz, then band-pass filtered (300-3000 Hz, second order Butterworth filters), from which the multiunit activity (MUA) was extracted as negative deflections in the electrical trace with an amplitude exceeding 4 SD of the baseline hash. Single units were separated from MUA using a wavelet-based spike sorting package (wave_clus) (Quiroga et al., 2004). Single unit isolation was confirmed based on the inter-spike-interval histogram (fewer than 3% of the spikes in the 0-3 ms bins) and the consistency of the spike waveform (SD of peak-to-trough delay of spikes within the cluster less than 0.2 ms).

### Optogenetic activation

#### Light delivery

Collimated blue light (488 nm) was generated by a diode laser (LuxX, Omicron) coupled to the implanted optic fiber assembly. Prior to implantation, laser power was recorded at the tip of each optic fiber assembly with a photodiode power sensor (Thorlabs, Inc.).

#### Evolutionary stimulus search procedure

The evolutionary stimulus search procedure used here follows the approach described previously for closed-loop optimization of visual and auditory stimulus parameters (Yamane et al., 2008; Chambers et al., 2014). Here, we apply this algorithm to the laser command signal rather than to the sensory stimulus. Voltage command signals to the laser were varied across five different parameters: pulse rate (5 to 50 Hz in 5 Hz increments), pulse width (5 to 19 ms in 2 ms increments), onset asynchrony between the laser and noise burst (−200 to 200 ms in 50 ms increments), peak power (0 to 30 mW in 5 mW increments) and duration (50 to 450 ms in 100 ms increments) yielding 25,200 unique permutations of laser parameters. Each run of the adaptive search procedure was initialized by randomly selecting 48 different laser settings. On every trial, laser stimulation was accompanied by a 250 ms, 40 dB SPL noise burst (4 ms raised cosine onset/offset ramps). Each trial was 2 s in duration.

Firing rate responses (the average of two repetitions) were calculated during the stimulus window (0-250 ms) and all stimuli played in the experiment thus far were rank-ordered at the end of each generation. At the conclusion of each evolutionary generation, the mean stimulus-evoked firing rate associated with each offspring were rank-ordered and the top ten most (or least) effective offspring were used as ‘breeders’ for the next generation. Offspring “evolved” from a parent breeding stimulus by randomly shifting one or more laser parameters to its nearest-neighbor value (for example, if the power of a breeder stimulus was 15 mW, its offspring could have a power of 10, 15 or 20 mW, as the sampling density for power was 5 mW). After the first generation, 37/50 stimuli were derived by the evolutionary algorithm, while 10/50 stimuli were chosen randomly from the entire acoustic feature space to avoid focusing on local maxima and mitigate potential decreases in firing rate response magnitude due to adaptation. The most (or least) effective stimulus from the first generation (the “yardstick” stimulus) was repeated in every subsequent generation to estimate the overall response stability across generations. Each generation featured 2/50 control stimuli consisting only of noise burst stimulus without laser stimulation.

Two criteria were used to safeguard against contamination of neural responses by movement, or other sources of noise. First, a global ceiling on firing rate was set to 400 Hz. Any stimuli corresponding to neural responses exceeding this ceiling were excluded from the dataset. Second, responses to stimuli across two repetitions were compared. If the neural responses to the two repetitions differed by more than 25Hz, at least one presentation was considered an artifact and the offspring was excluded from the dataset.

#### Stimulus effectiveness

To explicitly assess the effectiveness of the evolutionary design procedure, we devised a direct comparison of three conditions: *i)* sound alone, consisting of a 250ms white noise burst (40 dB SPL, 250 ms duration); *ii)* sound plus a “generic” activation of ACtx (250 ms, continuous 10 mW laser presented concurrently with the sound stimulus); *iii*) sound with “optimized” activation of ACtx, defined either as top- or bottom-performing offspring from the evolutionary search procedure. The inter-stimulus interval was 1 s and 50-100 trials were averaged for all three conditions.

#### One-dimensional tuning functions

One-dimensional tuning functions were obtained by varying one of the five laser parameters along their entire sampling range, while holding the remainder at their optimal value. Firing rates to all sound with laser trials were contrasted to interleaved trials in which sound was presented without laser. Average firing rates were computed from 10-20 repetitions of each unique condition.

### Electrophysiological data analyses

#### Frequency response areas

FRAs were delineated using pseudorandomly presented pure tones (50ms duration, 4ms raised cosine onset/offset ramps) of variable frequency (4-64 kHz in 0.1 octave increments) and level (0-60 dB SPL in 5 dB increments). Each pure tone was repeated two times and responses to each iteration were averaged. Spikes were collected from a 50 ms window beginning at stimulus onset. The tone-driven portion of the FRA was calculated using an automated method (Guo et al., 2012) and was used to determine the best frequency (BF; the frequency associated with the highest spike count, summed across all sound levels), and the bandwidth (measured 10dB above threshold). Recording sites along a single electrode shank were labelled as either central nucleus of the IC (CIC) or not central nucleus of the IC (NCIC), depending on the presence or absence of a tonotopic gradient as a function of depth (evaluated via a linear fit across BF’s from the recording sites on each shank). CNIC sites for a given shank were further categorized as broad or narrow (CICb and CICn, respectively) according to whether the tuning bandwidth measured 20 dB SPL above thresholds was > 1.75 octaves or < 1.75 octaves, respectively.

#### Selectivity asymmetry index

To quantify the directionality of the change in firing elicited by the laser, a selectivity asymmetry index was defined as

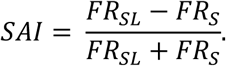

Here, *FR*_*sL*_ is the firing rate for the sound and laser condition and *FR*_*s*_ is the firing rate for the sound alone condition. The SAI is bounded between −1 and 1, with a value of 0 indicating equivalence between the two conditions and negative or positive values indicating either suppression or enhancement by the laser, respectively.

#### Quantification of one-dimensional tuning functions

To validate whether the evolutionary search algorithm was able to identify the true maximum, a measure of estimation error was evaluated from each one-dimensional tuning functions. This was defined as

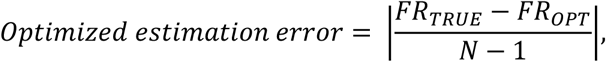

where *FR*_*TRUE*_ is the firing rate in response to the most efficient stimulus (the true maximum of the function), *FR*_*OPT*_ is the firing rate in response to the optimized stimulus, and *N* is the number of stimuli in the specific one-dimensional tuning function. The “random” error was computed by replacing *FR*_*OPT*_ with the mean firing rate from 1000 random draws from the one-dimensional function (excluding *FR*_*OPT*_).

To estimate the leverage of each stimulus dimension on the maximal neural response, a series of glyph plots were constructed. Each spoke in a glyph represents the z-score of the peak response relative to the distribution of all responses, where longer spokes indicate that a particular laser dimension had a disproportionately strong influence on a given neuron’s firing rate. In the rare cases where a neuron was lost prior to completion of the full stimulus protocol, no spoke was added for the missing dimensions.

To characterize the relative impact of each laser parameter on the overall variation in firing rates, a measure of firing rate leverage was quantified by computing the number of standard deviations between the peak of each one-dimensional tuning function and its mean (a z-score).

### Anatomy

Mice were deeply anesthetized with ketamine and transcardialy perfused with 4% paraformaldehyde in 0.01M phosphate buffered saline. The brains were extracted and stored in 4% paraformaldehyde for 12 hours before transferring to cryoprotectant (30% sucrose) for 48 hours. Sections (40µm) were cut using a cryostat (Leica CM3050S), mounted on glass slides and coverslipped (Vectashield). Fluorescence photomicrographs were obtained with a confocal microscope (Leica).

### Statistical analyses

All statistical analysis was performed with MATLAB (Mathworks). Descriptive statistics are reported as mean ± SEM, or median ± 95% confidence interval when data samples did not meet the assumptions of parametric statistical tests. In cases where the same data sample was used for multiple comparisons, we used the Holm-Bonferroni method to control for Type-I error inflation. Statistical significance was defined as p< 0.05.

## Results

To characterize the influence of cortical activation on midbrain sound processing, we injected a viral construct into the ACtx of adult mice to express Chronos, a channelrhodopsin with high sensitivity and the fastest channel kinetics of any opsin described to date **(Fig. 1a)** (Klapoetke et al., 2014; Guo et al., 2015). After allowing several weeks for the virus to incubate, we prepared mice for awake head-fixed recordings **(N** = 10). We made extracellular recordings of single units from all subdivisions of the IC using a 32-channel silicon probe **(Fig. 1b).** Post-mortem visualization of Chronos-EYFP in fixed tissue revealed dense labeling throughout all layers of Actx and a well-defined plexus of cortical axon terminals throughout the external and dorsal cortex of the IC (ECIC and DCIC, respectively) **(Fig. 1c).** A closer inspection revealed additional sparse labeling of corticocollicular axons innervating the central nucleus of the IC (CIC), as reported previously (Beyerl, 1978; Saldaña et al., 1996) (Fig. 1c, inset). Each shank on the recording probe sampled neural activity across a 0. 7 mm vertical expanse at 0.1 mm resolution **(Fig. 1d).** Units along a single shank were operationally assigned to being within or not within the central nucleus (CIC and NCIC, respectively) according to whether the best frequency (BF) increased tonotopically across recording depth. CIC units were further grouped as either broad/v-shaped or narrow/I-shaped according to the frequency tuning bandwidth measured 20 dB above threshold (CICb [> 1.75 octaves] and CICn [< 1.75 octaves], respectively).

**Figure 1.**
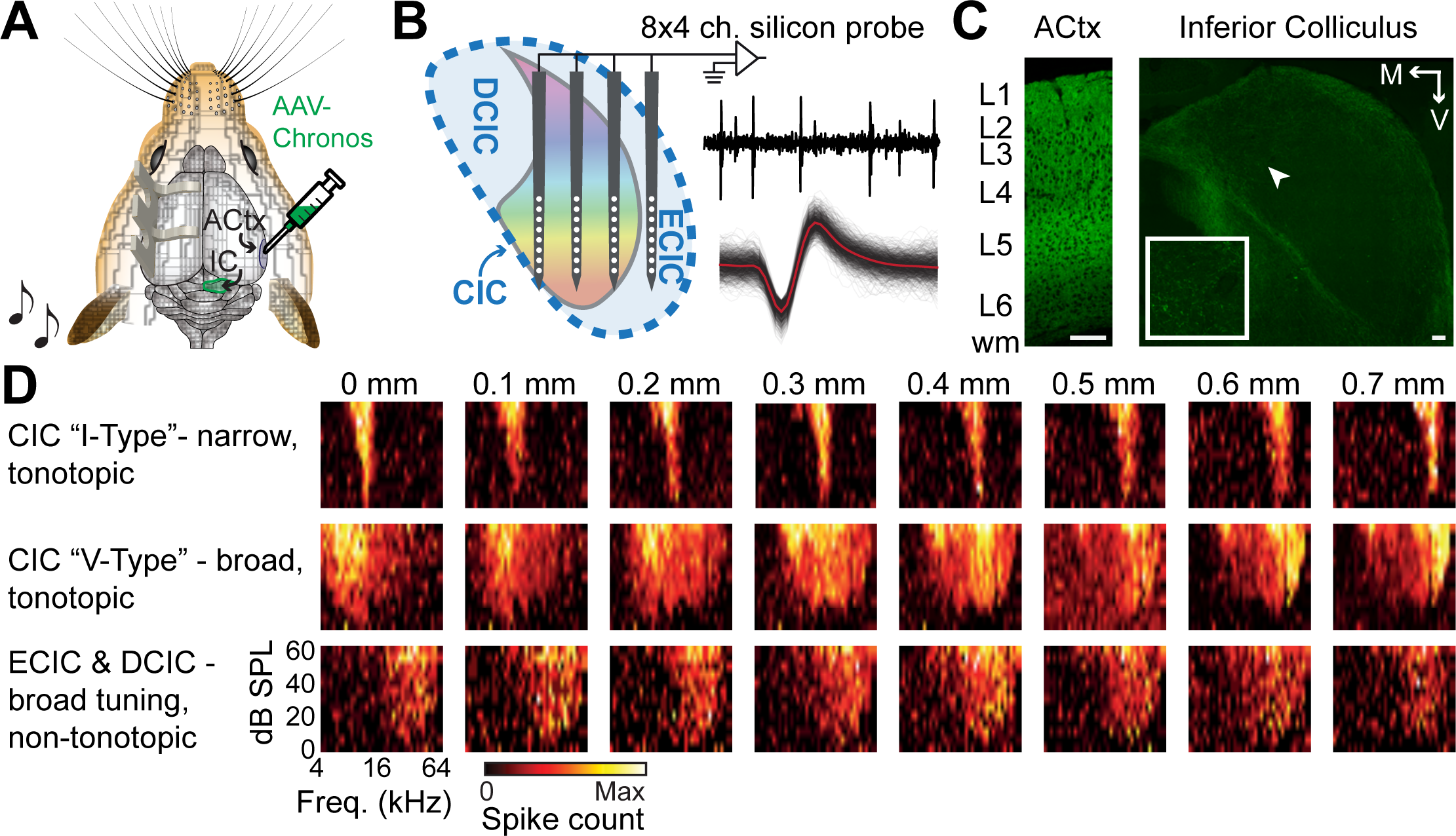
Functional parcellation of the mouse inferior colliculus. **(A)** Chronos, a fast channel rhodopsin, was expressed in the ACtx of adult mice with a viral construct. Weeks later, mice were prepared for awake, head-fixed IC recordings. **(B)** Single units were recorded across the medial-lateral and dorsal-ventral axes of the IC with 32-channel silicon probes with 200µm inter-shank separation and lO0µm spacing between individual contacts along a shank. A representative extracellular recording trace is shown alongside spike waveforms from an isolated single unit. (C) Chronos-EYFP expression is found in all layers of the ACtx (left) and in corticofugal axons terminating in the IC (right). CCol axons are predominantly clustered in the external cortex of the IC, but sparse terminal expression is also found in the central nucleus (white arrow, inset). Scale bars= 0.1 mm for both. L=layer, wm = white matter, M = medial, V=ventral. **(D)** Representative frequency response areas recorded from single units along individual shanks of the silicon probe.

### IC response modulation with concurrent cortical activation

Having operationally defined single units as CICn, CICb or NCIC, we then characterized the influence of cortical activation on sound-evoked single unit spiking. As a first pass, we presented a broadband noise burst to the contralateral ear and contrasted firing rates when sound was presented alone versus in combination with optogenetic stimulation via an implanted optic fiber resting atop the surface of the Actx **(Fig. 2a-b).** We probed the functional influence of the ACtx on IC units using a generic optogenetic stimulation protocol, in which a flash of moderately intense laser (10 mW) is presented concurrently with the stimulus.

**Figure 2.**
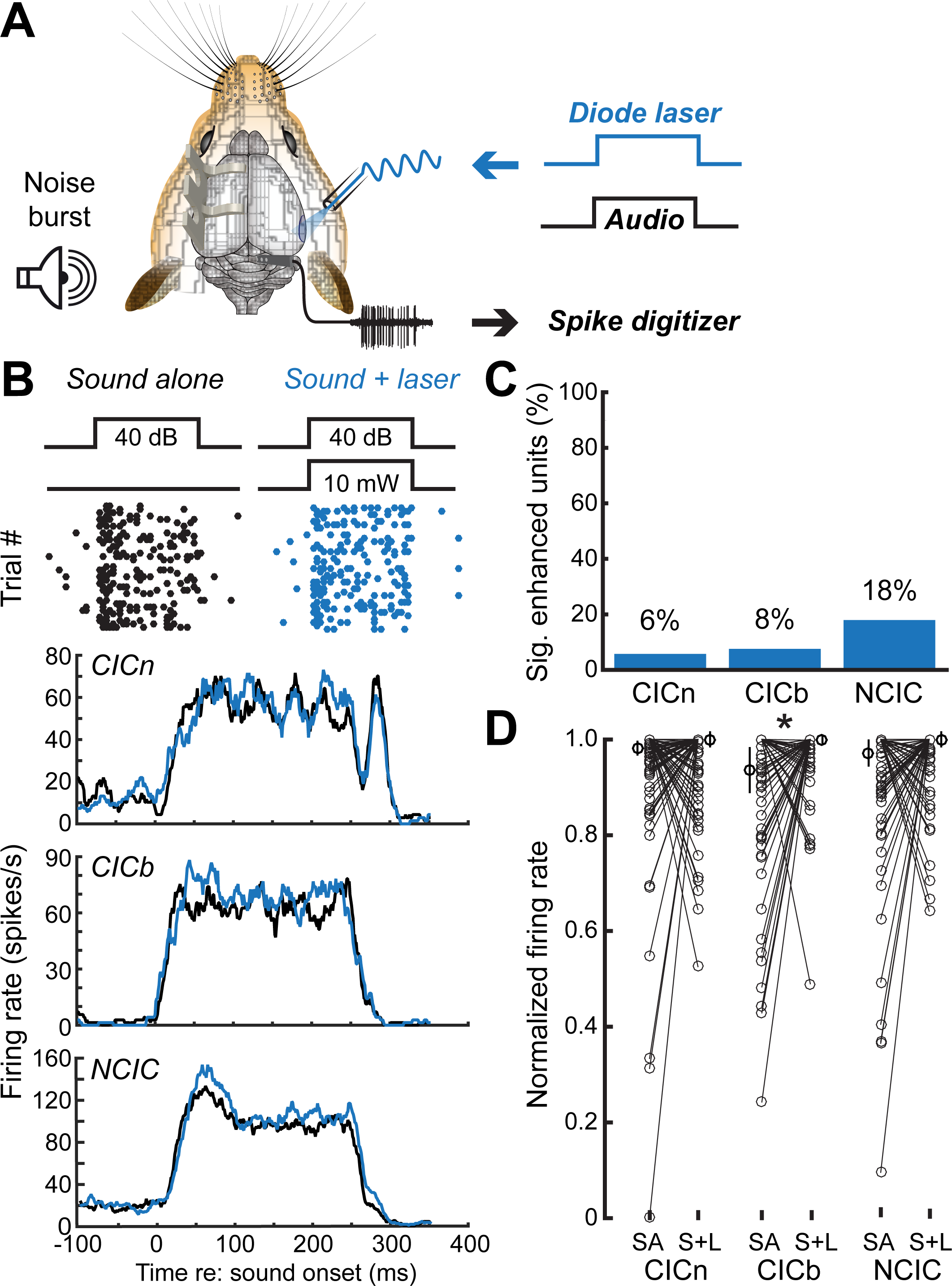
Modest enhancement of IC sound responses with concurrent ACtx activation. **(A)** Schematic of the paradigm to record sound-evoked IC single unit spiking while optogenetically activating ACtx with a generic laser pulse. **(B)** Spike rasters (top) and PSTHs (bottom) depict sound-evoked activity with and without concurrent ACtx activation from example single units from regions of the central nucleus with narrow and broad tuning (CICn and CICb, respectively) or regions outside of the central nucleus (NCIC). Percentage of units with significantly increased firing rates in the sound plus laser condition. **(D)** Normalized firing rates for sound alone versus sound plus laser trials (SA and S+L, respectively). Thin lines show individual units, values to either side depict the median and 95% confidence intervals. Asterisk reflects p < 0.05 with the Wilcoxon Sign Rank test.

Despite the dense network of cortical axons innervating the NCIC and, to a lesser extent, the CIC, the effect of cortical stimulation on downstream midbrain neurons was fairly modest. Sound-evoked firing rates were only significantly elevated in the sound + laser condition in < 15% of all recorded IC units **(Fig. 2c).** To measure the magnitude of corticofugal enhancement for each unit, we compute the normalized firing rate for each unit during the sound alone and sound + laser conditions. Cortical activation effects were weak and statistically not significant in the NCIC (n = 56) and CICn (n = 69; Wilcoxon signed-rank test, p = 0.11 and 0.65, respectively) but imposed a modest (6%) yet statistically significant increase in sound-evoked rates in CICb units (n = 53; p = 0.006, **Fig. 2d).**

### Converging on optimized optogenetic stimulation parameters with closed-loop spike feedback

The temporal patterning of cortical activation could be implemented in virtually limitless varieties. It therefore seemed premature to conclude that cortical activation has subtle effects on sound-evoked IC firing rates having only tried a single, generic form of activation. Given that the parameter space for optogenetic activation is vast and that cortical modulatory effects on IC neurons could be complex and non-linear, we reasoned that an exhaustive, brute force search of laser stimulation parameters on IC firing rates would not be feasible. Instead, we implemented a closed-loop search procedure designed to converge on maximally effective laser stimulation parameters based on single unit spike feedback. Optimization algorithms can often get stuck exploring local minima and maxima without ever exploring the regions of parameter space that elicit the most extreme firing rate variations. Genetic algorithms can avoid perseverating in local firing rate maxima by incorporating random “mutations” into each generation so that some resources are continually allocated towards exploring new regions of the stimulus manifold while others are invested in exploring local features in an identified effective region.

We repurposed a variation of a genetic algorithm that has been used to rapidly converge on complex shape stimuli to drive single units in visual cortex or spatial and spectrotemporal sound features to drive single units in the auditory cortex (Yamane et al., 2008; Chambers et al., 2014). Rather than use spike feedback to identify optimal acoustic stimulus parameters for a neuron, we kept the stimulus constant and used fluctuations in the firing rate of an individual IC neuron to iteratively tailor the voltage command signal to the laser. We varied five parameters of the optogenetic stimulation simultaneously: Pulse rate, the duration of the pulse train, the width of individual laser pulses, the amplitude (i.e., power) of each pulse, and the onset asynchrony between the noise burst and laser pulse train **(Fig. 3a).** We defined the minimum, maximum and reasonable sampling densities for each parameter, yielding 25,200 possible unique permutations of laser activation settings **(Fig. 3b).**

**Figure 3.**
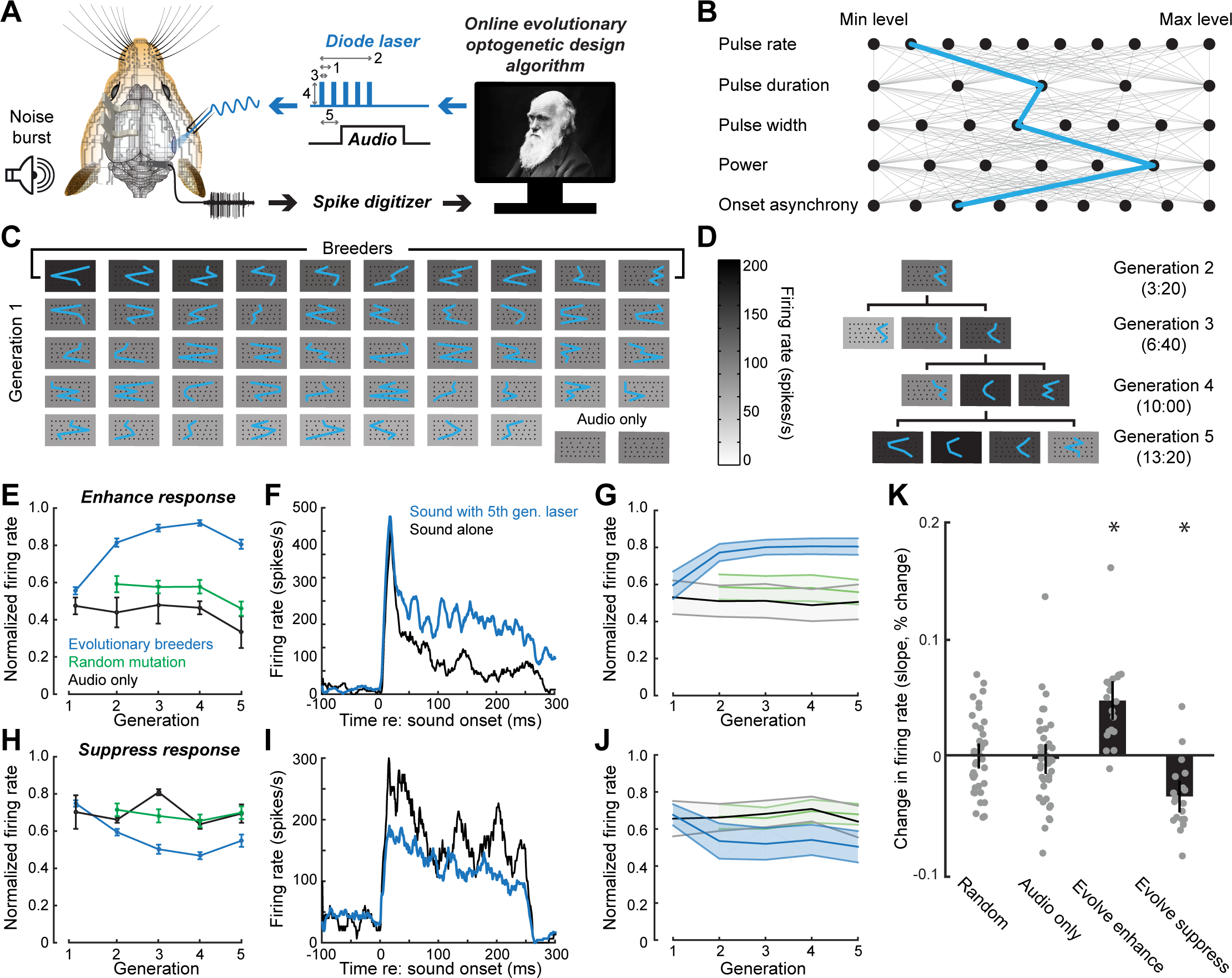
Optimizing optogenetic stimulation parameters based on closed-loop spike feedback. **(A)** Schematic of an evolutionary stimulus optimization procedure that uses variations in sound-evoked IC single unit spiking to configure the voltage command signal sent to the laser. The procedure specified five features (1-5): pulse rate, pulse duration, pulse width, power and onset asynchrony with respect to an invariant white noise burst, respectively. **(B)** The five laser parameters were tested across a pre-determined range, yielding 25,200 unique combinations, an example of which is shown in blue. (C) Example responses of one IC unit to 48 randomly chosen cortical activation configurations, where the grayscale background represents the firing rate during the 250 ms white noise burst. Blue lines depict the laser settings for each stimulus, borrowing from the plotting convention illustrated in *B.* IC firing rates are rank-ordered by firing rate, where the top 10 are selected as “breeding” stimuli for the next generation. Two trials per generation are audio only controls. **(D)** Example of a breeder stimulus selected from Generation 2 that evolved to elicit higher firing rates in subsequent generations (bottom 3 rows) by adjusting features to nearest neighbor values. Elapsed time since the start of the evolutionary search process is shown for each generation. (E) Example unit showing changes in the mean ± SEM normalized firing rate for experiments where the algorithm was directed to enhance IC sound responses. Mean SEM firing rate for the top 10 stimuli of each generation selected as breeders, the 10 stimuli chosen at random and the 2 audio only control stimuli. **(F)** Example unit PSTH associated by a 250 ms noise burst presented alone (black) or with a breeder optogenetic stimulation pattern identified in the 5^th^ generation of the stimulus search. **(G)** Mean ± SEM change in normalized firing rate across the sample of units (n = 21). **(H-J)** Same as f-G, but in unit recordings where the algorithm was directed to suppress IC sound responses (n = 20). **(K)** Change in normalized firin g rate for breeding stimuli, random stim uli and audio only control stimuli were quantified by calculatin g the slopes of linear fits. IC unit responsesto noise bursts presented alone or with randomly selected optogenetic stimuli were unchanged over the course of testing, while responses to the most effective breeding stimuli were significantly enhanced or suppressed, corresponding to whether the algorithm was instructed to increase or decrease IC firing rates with ACtx activation. Data are medians± 95% confidence interval. Asterisks represent significant differences using a one-sample signed rank tests against a population mean of zero.

### Bi-directional control over IC spike rates with optimized cortical stimulation

Once an IC single unit was isolated, we interrogated it with 48 randomly selected combinations of laser activation parameters and 2 control conditions consisting of noise bursts without cortical activation **(Fig. 3c).** The 48 sound and laser combinations were rank-ordered by firing rate and the top 10 most effective settings were identified as “breeders” that would constrain the properties of the subsequent generation. “Offspring” for an individual breeder were created by randomly shifting one or more laser parameters to its nearest-neighbor value; for example, if the laser pulse rate was 10Hz, it might be shifted to 5 or 15 Hz. In so doing, we could identify a lineage of increasingly “fit” cortical activation settings that began with a randomly selected breeder in generation 1 that produced offspring of increasing-but varying-effectiveness across five generations **(Fig. 3d).** Following the initial generation of random laser patterns, each subsequent generation included 37 offspring that were defined by the evolutionary design algorithm, 10 conditions selected at random, 2 audio only controls and a “yardstick” stimulus, defined as the most effective condition from Generation 1. Whereas IC firing rates to the audio alone control or to noise bursts paired with randomly selected laser patterns were constant across generations, the evolutionary search procedure could converge on increasingly effective activation parameters in just a few minutes **(Fig. 3e-g).**

Although corticocollicular projections are glutamatergic (Kaneko et al., 1987; Feliciano and Potashner, 1995), we reasoned that it might be possible for the evolutionary procedure to identify cortical activation patterns that suppressed midbrain spiking if, for example, there were a particular temporal patterning of cortical inputs that disproportionately activated inhibitory interneurons within the IC. We tested this idea by programming the evolutionary algorithm to identify the least effective laser activation parameters and breed offspring to converge on conditions associated with the weakest sound-evoked firing rates. We found that cortical activation could also suppress IC firing rates relative to sound-alone controls, where firing rates to the least effective breeder stimuli became progressively lower across generations **(Fig. 3h-j).** To quantify these observations, we linearly fit the normalized firing functions across generations for audio only controls, randomly selected laser parameters, and patterns identified through the evolutionary search procedure as optimally enhancing or optimally suppressing **(Fig. 3k).** We found that firing rate slopes were significantly increased across generations when the algorithm was set to enhance (n = 21, p = 0.00009, one-sample Wilcoxon Signed-Rank test against a population mean of zero), were significantly decreased when the algorithm was set to suppress (n = 20, p = 0.03), but were not significantly changed for control conditions where sound was presented without laser or where sound was presented with laser settings selected at random (n =41 for each, p = 0.4 and 0.76, respectively).

### Distributed effects of optimized cortical stimulation throughout the IC

Once a maximally enhancing or suppressing optogenetic stimulation protocol was identified through the evolutionary design process, we could then compare this “optimized” activation parameter to other conditions in single units identified through offline spike sorting **(Fig. 4a).** The evolutionary design algorithm used the firing rate of just one single unit to steer the stimulus design procedure. Having established that the stimulus design process was effective at enhancing or suppressing the firing rate of this “driver” unit (Fig. 3k), we next addressed whether this pattern was associated with commensurate firing rate modulation in other IC “passenger” units that were along for the ride **(Fig. 4b).**

**Figure 4.**
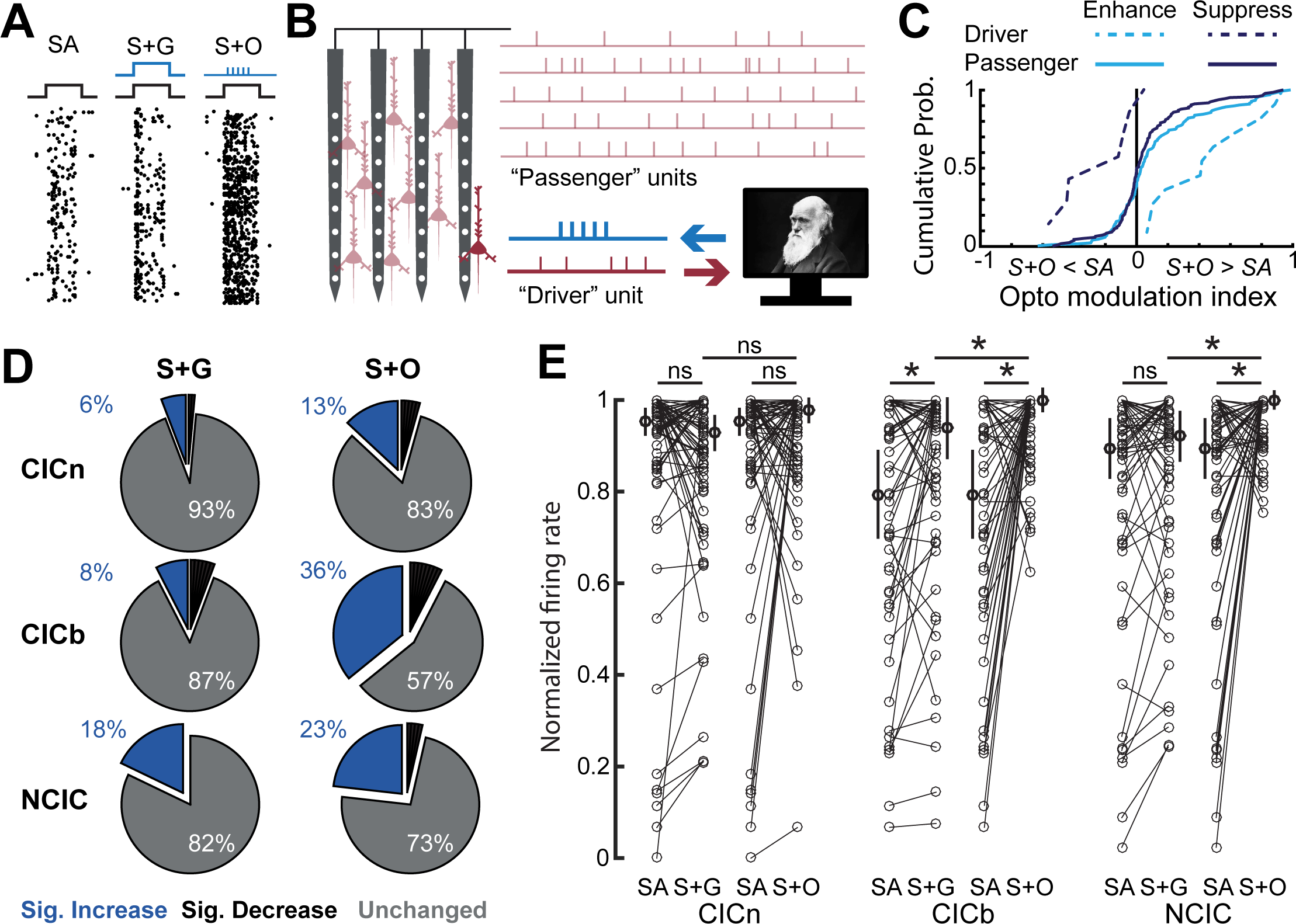
ACtx activation has stronger effects on IC firing rates with optimized optogenetic stimulation parameters. **(A)** Spike rasters from an example IC unit in response to a 250 ms sound alone (SA, left), sound with a generic concurrent activation of ACtx (S+ G, middle) and sound presented with an optogenetic laser pulse optimized through the evolutionary search procedure (S+O, right). **(B)** Cartoon illustrating that in addition to the single unit that drives the evolutionary search algorithm, there are many other “passenger” units recorded simultaneously on other probe contacts. **(C)** The distribution of optogenetic modulation values from all IC driver and passenger units, according to the formula (S+O-SA) / (S+O + SA), where zero (vertical black line) indicates an equivalent firing rate between SA and S+O. **(D)** Percentage of passenger recordings sites with significantly modulated firing rates for each functional classification of IC unit type. Data from the S+O condition reflect recording blocks when the evolutionary design procedure was programmed to enhance firing rates. **(E)** Change in normalized firing rate with generic versus optimized ACtx stimulation. Thin lines show individual units, values to either side depict the median and 95% confidence intervals. Asterisks reflect p < 0.05 with the Wilcoxon Sign Rank test, after correcting for multiple comparisons with Holm-Bonferroni.

As a first step, we contrasted IC firing rates evoked by the sound alone versus sound plus optimized cortical activation using an asymmetry index bounded between −1 and 1, where positive values reflected higher firing rates to sound plus optimized, negative values reflected higher firing rates to sound alone, and equivalent firing rates would yield a value of zero **(Fig. 4c).** In cases where the same single units that drove the evolutionary procedure could be identified and held for additional recordings, we confirmed that spiking for these driver units were significantly suppressed when the algorithm was programmed to suppress firing and were significantly enhanced when the algorithm was programmed to enhance firing rates (One-sample Wilcoxon signed-rank test against a population mean of zero, Enhance drivers, n =11, p = 0.001, Fig. 4d light dashed line; Suppress drivers, n = 7, p = 0.03, Fig. 4d dark dashed line). Passengers tested with an optogenetic activation pattern tailored for another unit went along for the ride by also exhibiting significantly enhanced firing rates to the optimized activation condition (n = 168, p = 0.0001, Fig. 4d light solid line), but we did not observe any differences in IC passenger units when the driver was suppressed (n = 149, p = 0.06; Fig. 4c dark solid line). These findings suggest that while cortical activation can be patterned in a manner that suppresses sound-evoked midbrain firing rates, these effects were spatially localized such that neighboring units were unaffected or may have even increased their firing rates, perhaps providing the basis for local spiking suppression (Yang et al., 1992; Schofield and Beebe, 2018). Increasing IC firing rates with optimized activation of cortical neurons, by contrast, reflected a more widespread enhancement that could also be observed in passenger units that did not guide the optogenetic stimulus design. All subsequent analyses focus only on firing rate changes in passenger units with Actx activation protocols that were designed to enhance firing.

To more closely investigate how optimized cortical activation affected the firing rates of nearby IC passenger units, we returned to the functional trichotomy of IC response types (CICn, CICb and NCIC) and contrasted responses to sound presented with and without the optimized activation waveform to sound presented with the generic laser waveform described in in Fig. 2. Compared to the small minority of IC units that were significantly enhanced by a generic cortical activation waveform, using an optimized cortical activation waveform more than doubled the percentage of significantly increased IC units **(Fig. 4d).** With that said, narrowly tuned units in the CIC remained the least sensitive to cortical activation; even with an optimized activation pattern, only 13% were significantly enhanced and magnitude of firing rate change was not significantly different overall than sound alone, nor different than sound paired with generic cortical activation (n = 64, WSR test corrected for multiple comparisons, S+O vs SA, p = 0.98; S+O vs S=G, p = 0.22; **Fig. 4e, left).** By contrast, CICb units showed significantly enhanced sound-evoked responses with optimized pattern of cortical activation, both compared to sound alone (n = 59, WSR, p = 0.0002) and to sound presented with generic activation waveform (WSR, p = 0.02; **Fig. 4e, middle).** NCIC units were also significantly more responsive to sound paired with the optimized activation pattern than to sound alone (n = 56, WSR, p = 0.002) and were also significantly more responsive to sound presented with the optimized activation pattern than to sound presented with generic activation (WSR, p = 0.003 **Fig. 4e, right).** Differences between generic and optimized patterns of cortical activation were consistent across mice where the optic fiber was implanted atop the ACtx - and therefore indiscriminately activated all infected neurons-versus mice where the fiber was implanted between the thalamus and the midbrain, to selectively activate fasciculated corticofugal axon bundles entering the tectum (Wilcoxon Rank Sum statistic on specific versus nonspecific stimulation, p > 0.63 for CICb, CICn, and DCIC). Data reported throughout therefore were combined across mice with fiber tips in both locations.

### Identifying the most influential parameters for optimized cortical stimulation

Because optimized search algorithms test only a portion of the stimulus manifold, there is always some doubt as to whether the optimal stimulus identified by the algorithm is the absolute maximum. We addressed this concern by allowing 20% of the laser configurations in generations 2-5 to be selected from random positions in the stimulus manifold. Still, the possibility remained that the optimization procedure might have perseverated on a local maximum adjacent to the true peak or might have been led astray by spurious responses on outlying trials. To validate the evolutionary design approach described here, a subset of neurons were subjected to a subsequent test in which a single laser parameter was varied while the other four were held at the optimal value identified by the evolutionary search procedure. IC units could show saturating, monotonically increasing or non-monotonically tuned responses as a given laser parameter was varied from the minimum to maximum of its range **(Fig. 5a).**

**Figure 5.**
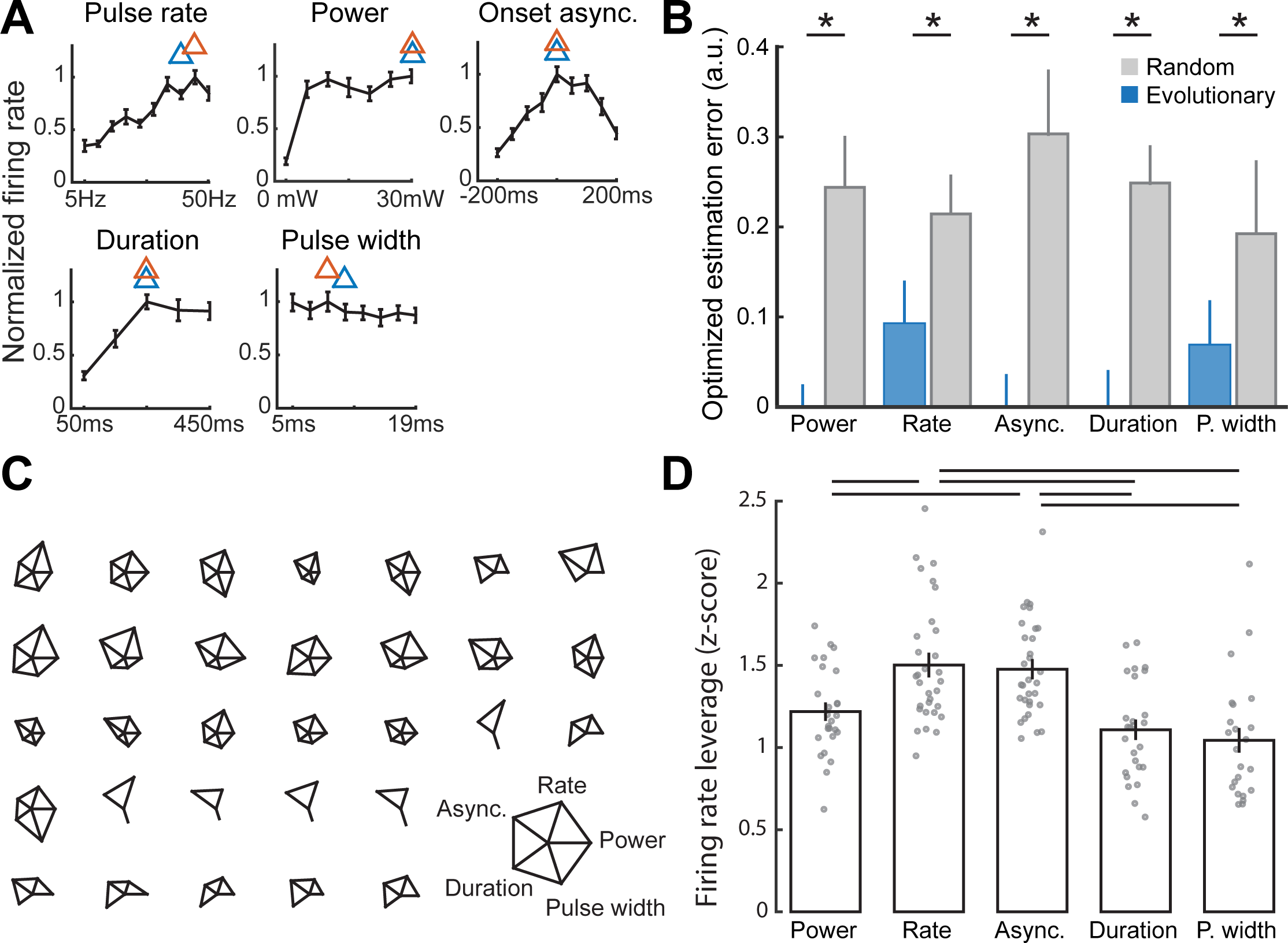
Evolutionary design procedure rapidly converges on highly effective regions of the laser command signal manifold. **(A)** One-dimensional tuning functions from an example IC unit where each dimension of the optimized activation stimulus was systematically varied while the other four were held constant. Orange and blue triangles indicate the actual peak and the optimal value suggested from the evolutionary search procedure, respectively. **(B)** Estimation error between the true maximum and optimized value identified through the evolutionary procedure or selected at random. Data are medians ± 95% confidence interval. Asterisks represent p < 0.05 with a Wilcoxon Rank Sum test. **(C)** Glyph plots constructed from all IC units subjected to one-dimensional variations of the preferred stimulus. Each represents the difference in firing rate between the maximum and minimum values of each one-dimensional tuning function, meant to estimate the leverage of each stimulus dimension on the maximal response of the neuron. **(D)** Leverage of each individual laser parameter on the overall variation in firing rates. Individual units are shown as gray symbols. Data are medians ± 95% confidence interval. Horizontal lines reflect p < 0.05 with the Wilcoxon Sign Rank test, after correcting for multiple comparisons with Holm-Bonferroni.

We observed that the optimal peak activation parameters identified during the evolutionary search procedure was a close match to peak response identified through this exhaustive one-dimensional search procedure. We computed the ordinal difference in stimulus position between the true peak and the best stimulus identified by the evolutionary search procedure and compared the estimation error to what would have occurred by chance **(Fig. 5b).** We found that the evolutionary search procedure converged on or near the true optimal value with significantly less error that what would have occurred by chance alone for all five laser parameters (Power, n = 26, Wilcoxon sign-rank test, p = 0.0001; Rate, n = 31, p = 0.0003; Asynchrony, n = 31, p < 0.00001; Duration, n = 26, p < 0.00001; Pulse width, n = 23, p = 0.02). Glyph plots were constructed for all units where each spoke represents the Z-score of the peak response relative to the distribution of all responses, where longer spokes indicate that a particular laser dimension had a disproportionately strong influence on IC firing rates **(Fig. 5c).** Comparing across optogenetic activation parameters, we found that the laser pulse rate and laser-sound onset asynchrony exerted the greatest overall leverage on sound-evoked IC firing rates **(Fig. 5d).**

## Discussion

We first identified recording sites in the dorsal and external cortex of the IC and functionally parceled regions of the CIC according to frequency tuning bandwidth (Fig. 1). We reported that optogenetic activation of Actx neurons induced only a 5-10% change in sound-evoked IC firing rates (Fig. 2). The percentage of significantly modulated units was highest in the NCIC, as expected from the relative abundance of CCol axons in the external and dorsal cortex. Interestingly, the cortical modulation effects were largest in magnitude in broadly tuned cells within the CIC, suggesting that broadly-tuned CIC units may integrate across a wider range of synaptic inputs, whether those are bottom-up inputs evoked by tones or descending inputs elicited by optogenetic activation (Chen et al., 2018). We then used a closed-loop optimization strategy to readily converge on patterns of cortical activation that were able to enhance or suppress the firing rate of a chosen downstream IC neuron (Fig. 3). Given that corticocollicular projections are glutamatergic (Kaneko et al., 1987; Feliciano and Potashner, 1995), the suppression we observed is likely due to di-synaptic activation of inhibitory interneurons with the IC (Schofield and Beebe, 2018). In cases where the closed-loop optimization enhanced an IC neuron, neighboring neurons recorded simultaneously were also enhanced, and this enhancement was again most pronounced in CICb (Fig. 4). Conversely, in cases where the closed-loop optimization suppressed an IC neuron, neighboring units typically did not change their firing rates. This indicates that suppressive patterns of cortical activation led to more spatially localized collicular effects, likely through interactions with the local inhibitory circuitry (Ito et al., 2016). Finally, we identified that pulse rate and laser-sound onset asynchrony were able to best leverage changes in sound-evoked IC firing rates (Fig. 5).

Closed-loop algorithms for stimulus optimization are often used in sensory neurophysiology for firing rate control, where a sensory stimulus is optimized such that the firing rate of a chosen neuron is maximized (Bleeck et al., 2003; O’Connor et al., 2005; Yamane et al., 2008; Hung et al., 2012; Koelling and Nykamp, 2012; Chambers et al., 2014). Here, we modified a genetic algorithm previously used to optimize sound features, and tasked it with optimizing patterns of ACtx activation in a closed-loop fashion (Chambers et al., 2014). Genetic algorithms are particularly suited to our purpose because they are not “local search” methods and, as a result, are particularly robust to local maxima and more extensively sample the stimulus space (DiMattina and Zhang, 2013).

Although closed-loop feedback has previously been used to optimize both optical and electrical stimulation, its use has been more focused on real-time instantaneous feedback control, rather than stimulus optimization (Wagenaar, 2005; Wallach et al., 2011; Newman et al., 2013, 2015). Moving beyond sensory characterization, closed-loop feedback has also been shown to improve brain-computer interfaces (Cunningham et al., 2011; Shanechi et al., 2016), to induce motor plasticity (Jackson et al., 2006), and to provide all-optical control of neural circuits (Zhang et al., 2018). Closed-loop firing rate control also has a direct translational relevance, where feedback could be used to guide sensory neural prosthetics, such as cochlear or retinal implants. Indeed, closed-loop control of deep brain stimulation has been used to improve therapies for Parkinson’s disease (Feng et al., 2007a, 2007b).

Our findings demonstrate that the feedback influence ACtx has on the IC can vary both in sign and degree dependent on how pre-synaptic ACtx neurons are activated in time. However, corticocollicular neurons do not exclusively synapse onto IC neurons; they collateralize onto multiple downstream structures including the thalamus and striatum (Asokan et al., 2018). Different ACtx activation patterns may impact these various structures differently, as downstream synaptic properties could lead to postsynaptic variation in response. Thus, it may be the case that the same set of presynaptic ACtx neurons could modulate neural firing of disparate downstream targets in different ways dependent upon the particular pattern of temporal activation. Similar effects have been observed in the functional influence of entorhinal cortex neurons on distinct downstream regions of CA1 (Igarashi et al., 2014). Future work could focus on whether the optimized optogenetic activation settings described might be naturally employed by ACtx neurons under listening conditions that would place a premium on dampening or enhancing ascending auditory activity (Guo et al., 2017).

## Acknowledgements

We thank Ed Boyden and Nathan Klapoetke for generously sharing the Chronos viral construct. This work was supported by NIH grants DC017078 (DBP), DC015376 (RSW) and the Bertarelli Fellowship in Translational Neuroscience and Neuroengineering (CV). DBP and KEH designed the experiments. KEH developed software control. CV and RSW collected all data. CV and RSW perfo rm ed data analysis. DBP and RSW wrote the manuscript.

## References

Asokan MM, Williamson RS, Hancock KE, Polley DB (2018) Sensory overamplification in layer 5 auditory corticofugal projection neurons following cochlear nerve synaptic damage. Nat Commun 9:1–10.

Beyerl BD (1978) Afferent projections to the central nucleus of the inferior colliculus in the rat. Brain Res 145:209–223.

Bleeck S, Patterson RD, Winter IM (2003) Using genetic algorithms to find the most effective stimulus for sensory neurons. J Neurosci Methods 125:73–82.

Chambers AR, Hancock KE, Maison SF, Liberman MC, Polley DB (2012) Sound-evoked olivocochlear activation in unanesthetized mice. J Assoc Res Otolaryngol 13:209–217.

Chambers AR, Hancock KE, Sen K, Polley DB (2014) Online stimulus optimization rapidly reveals multidimensional selectivity in auditory cortical neurons. J Neurosci 34:8963–8975.

Chen C, Cheng M, Ito T, Song S (2018) Neuronal Organization in the Inferior Colliculus Revisited with Cell-Type-Dependent Monosynaptic Tracing. J Neurosci 38:3318–3332.

Coomes DL, Schofield RM, Schofield BR (2005) Unilateral and bilateral projections from cortical cells to the inferior colliculus in guinea pigs. Brain Res 1042:62–72.

Cunningham JP, Nuyujukian P, Gilja V, Chestek CA, Ryu SI, Shenoy K V. (2011) A closed-loop human simulator for investigating the role of feedback control in brain-machine interfaces. J Neurophysiol 105:1932–1949.

Diamond IT, Jones EG, Powell TPS (1969) The projection of the auditory cortex upon the diencephalon and brain stem in the cat. Brain Res 15:305–340.

DiMattina C, Zhang K (2013) Adaptive stimulus optimization for sensory systems neuroscience. Front Neural Circuits 7:1–16.

Feliciano M, Potashner SJ (1995) Evidence for a Glutamatergic Pathway from the Guinea Pig Auditory Cortex to the Inferior Colliculus. J Neurochem 65:1348–1357.

Feng X-J, Shea-Brown E, Greenwald B, Kosut R, Rabitz H (2007a) Optimal deep brain stimulation of the subthalamic nucleus—a computational study. J Comput Neurosci.

Feng XJ, Greenwald B, Rabitz H, Shea-Brown E, Kosut R (2007b) Toward closed-loop optimization of deep brain stimulation for Parkinson’s disease: Concepts and lessons from a computational model. J Neural Eng 4.

Guo W, Chambers AR, Darrow KN, Hancock KE, Shinn-Cunningham BG, Polley DB (2012) Robustness of cortical topography across fields, laminae, anesthetic states, and neurophysiological signal types. J Neurosci 32:9159–9172.

Guo W, Clause AR, Barth-Maron A, Polley DB (2017) A corticothalamic circuit for dynamic switching between feature detection and discrimination. Neuron 95:180–194.e5.

Guo W, Hight AE, Chen JX, Klapoetke NC, Hancock KE, Shinn-Cunningham BG, Boyden ES, Lee DJ, Polley DB (2015) Hearing the light: neural and perceptual encoding of optogenetic stimulation in the central auditory pathway. Sci Rep 5:10319.

Hung CC, Carlson ET, Connor CE (2012) Medial Axis Shape Coding in Macaque Inferotemporal Cortex. Neuron 74:1099–1113.

Igarashi KM, Lu L, Colgin LL, Moser MB, Moser EI (2014) Coordination of entorhinal-hippocampal ensemble activity during associative learning. Nature 510:143–147.

Ito T, Bishop DC, Oliver DL (2016) Functional organization of the local circuit in the inferior colliculus. Anat Sci Int 91:22–34.

Jackson A, Mavoori J, Fetz EE (2006) Long-term motor cortex plasticity induced by an electronic neural implant. Nature 444:56–60.

Kaneko T, Urade Y, Watanabe Y, Mizuno N (1987) Production, characterization, and immunohistochemical application of monoclonal antibodies to glutaminase purified from rat brain. J Neurosci 7:302–309.

Klapoetke NC et al. (2014) Independent optical excitation of distinct neural populations. Nat Methods 11:338–346.

Koelling ME, Nykamp DQ (2012) Searching for optimal stimuli: Ascending a neuron’s response function. J Comput Neurosci 33:449–473.

Ludwig KA, Miriani RM, Langhals NB, Joseph MD, Anderson DJ, Kipke DR (2009) Using a Common Average Reference to Improve Cortical Neuron Recordings From Microelectrode Arrays. J Neurophysiol 101:1679–1689.

Ma X, Suga N (2001a) Plasticity of bat’s central auditory system evoked by focal electric stimulation of auditory and/or somatosensory cortices. J Neurophysiol 85:1078–1087.

Ma X, Suga N (2001b) Corticofugal modulation of duration-tuned neurons in the midbrain auditory nucleus in bats. Proc Natl Acad Sci 98:14060–14065.

Nakamoto KT, Jones SJ, Palmer AR (2008) Descending projections from auditory cortex modulate sensitivity in the midbrain to cues for spatial position. J Neurophysiol 99:2347–2356.

Newman JP, Fong MF, Millard DC, Whitmire CJ, Stanley GB, Potter SM (2015) Optogenetic feedback control of neural activity. Elife 4:1–24.

Newman JP, Zeller-Townson R, Fong M-F, Arcot Desai S, Gross RE, Potter SM (2013) Closed-Loop, Multichannel Experimentation Using the Open-Source NeuroRighter Electrophysiology Platform. Front Neural Circuits 6:1–18.

O’Connor KN, Petkov CI, Sutter ML (2005) Adaptive Stimulus Optimization for Auditory Cortical Neurons. J Neurophysiol 94:4051–4067.

Quiroga RQ, Nadasdy Z, Ben-Shaul Y (2004) Unsupervised spike detection and sorting with wavelets and superparamagnetic clustering. Neural Comput 16:1661–1687.

Robinson BL, Harper NS, McAlpine D (2016) Meta-adaptation in the auditory midbrain under cortical influence. Nat Commun 7:1–8.

Saldaña E, Feliciano M, Mugnaini E (1996) Distribution of descending projections from primary auditory neocortex to inferior colliculus mimics the topography of intracollicular projections. J Comp Neurol 371:15–40.

Schofield BR, Beebe NL (2018) Subtypes of GABAergic Cells in the Inferior Colliculus. Hear Res:1–10.

Shanechi MM, Orsborn AL, Carmena JM (2016) Robust Brain-Machine Interface Design Using Optimal Feedback Control Modeling and Adaptive Point Process Filtering. PLoS Comput Biol 12:1–29.

Wagenaar DA (2005) Controlling Bursting in Cortical Cultures with Closed-Loop Multi-Electrode Stimulation. J Neurosci 25:680–688.

Wallach A, Eytan D, Gal A, Zrenner C, Marom S (2011) Neuronal Response Clamp. Front Neuroeng 4:1–10.

Winer JA (2006) Decoding the auditory corticofugal systems. Hear Res 212:1–8.

Winer JA, Larue DT, Diehl JJ, Hefti BJ (1998) Auditory cortical projections to the cat inferior colliculus. J Comp Neurol 400:147–74.

Yamane Y, Carlson ET, Bowman KC, Wang Z, Connor CE (2008) A neural code for three-dimensional object shape in macaque inferotemporal cortex. Nat Neurosci 11:1352–1360.

Yan J, Ehret G (2002) Corticofugal modulation of midbrain sound processing in the house mouse. Eur J Neurosci 16:119–128.

Yan J, Zhang Y (2005) Sound-guided shaping of the receptive field in the mouse auditory cortex by basal forebrain activation. Eur J Neurosci 21:563–576.

Yang LC, Pollak GD, Resler C (1992) GABAergic Circuits Sharpen Tuning Curves and Modify Response Properties in the Moustache Bat Inferior Colliculus. J Neurophysiol 68:1760–1774.

Zhang Z, Russell LE, Packer AM, Gauld OM, Häusser M (2018) Closed-loop all-optical interrogation of neural circuits in vivo. Nat Methods.

Zhou X, Jen PHS (2005) Corticofugal modulation of directional sensitivity in the midbrain of the big brown bat, Eptesicus fuscus. Hear Res 203:201–215.

